# Plasmodesmata-dependent intercellular movement of bacterial effectors

**DOI:** 10.1101/2020.12.10.420240

**Authors:** Zhongpeng Li, Haris Variz, Yani Chen, Su-Ling Liu, Kyaw Aung

## Abstract

Pathogenic microorganisms deliver protein effectors into host cells to suppress host immune responses. Recent findings reveal that phytopathogens manipulate the function of plant cell-to-cell communication channels plasmodesmata (PD) to promote diseases. Several bacterial and filamentous pathogen effectors have been shown to regulate PD in their host cells. A few effectors of filamentous pathogens have been reported to move from the infected cells to neighboring plant cells through PD; however, it is unclear whether bacterial effectors can traffic through PD in plants. In this study, we systemically determined the intercellular movement of *Pseudomonas syringae* pv. *tomato* (*Pst*) DC3000 effectors between adjoining plant cells in *Nicotiana benthamiana*. We observed that at least 16 *Pst* DC3000 effectors move from transformed cells to the surrounding plant cells. The movement of the effectors is largely dependent on their molecular weights. The expression of PD regulators, Arabidopsis PD-located protein PDLP5 and PDLP7, lead to PD closure and inhibits the PD-dependent movement of a bacterial effector in *N. benthamiana*. Similarly, a 22-amino acid peptide of bacterial flagellin (flg22) treatment induces PD closure and suppresses the movement of a bacterial effector in *N. benthamiana*. Together, our findings demonstrated that bacterial effectors are able to move intercellularly through PD in plants.

## INTRODUCTION

Plasmodesmata (PD) are membrane-lined channels which physically connect adjoining plant cells. PD provide the symplastic pathway for the connected cells to exchange molecules directly and selectively (Lucas et al., 2009; Brunkard and Zambryski, 2017; Nicolas et al., 2017). The PD-dependent movement of hormones, sugars, proteins, and RNAs has been well documented in plants (Kragler, 2013; Schulz, 2015; Kitagawa and Jackson, 2017; Reagan et al., 2018). In addition to their fundamental roles in plant growth and development, recent findings highlighted the crucial roles of PD in plant immunity (Cheval and Faulkner, 2018).

PD enable the continuity of the plasma membrane (PM) and endoplasmic reticulum (ER) and link the cytoplasm of adjoining plant cells. The space between the PM and ER membrane lining, known as cytoplasmic sleeve, allows the trafficking of molecules between the adjoining plant cells. The function of PD is largely defined by their aperture in permitting molecules to move across. The largest molecules can traffic through the cytoplasmic sleeve is known as the size exclusion limit (Kim and Zambryski, 2005). Soluble green fluorescent proteins (1×sGFP; 27 kDa) can freely move between plant cells through PD, whereas the movement of 2×sGFP (54 kDa) and 3×sGFP (71 kDa) are largely inhibited between Arabidopsis cells (Kim et al., 2005; Aung et al., 2020). Among different regulators, callose plays the most prominent role in regulating the PD function. Callose is plant polysaccharide, which could be deposited in the cell wall around the PM lining of PD. The accumulation and degradation of callose at PD allow plant cells to dynamically control the closing and opening of PD. Callose accumulation at PD is positively correlated with the PD closure. Callose synthase (CalS) and ß-1,3-glucanase are involved in callose biosynthesis and degradation, respectively (De Storme and Geelen, 2014; Wu et al., 2018).

In addition to the enzymes directly involved in regulating callose homeosis at PD, plasmodesmata-located proteins (PDLPs) play critical roles in modulating the plasmodesmal function. Although the molecular mechanism of PDLPs is unknown, they affect callose homeostasis at PD. Ectopic expression of PDLP5 results in overaccumulation of callose, whereas a *pdlp5* knock-out mutant accumulates much less callose at PD compared to that of wild type in Arabidopsis (Lee et al., 2011). Different members of PDLPs have been demonstrated their roles in plant immunity. The expression of *PDLP5* transcripts is upregulated by *Pseudomonas syringae* pv. *maculicola* ES4326 infection (Lee et al., 2008; Lee et al., 2011). PDLP1 accumulates at the PM and haustorial interfaces during *Hyaloperonospora arabidopsidis* (*Hpa*) infection (Caillaud et al., 2014). The polar localization of PDLP1 at the haustorium leads to callose deposition at the interface (Caillaud et al., 2014). Despite the involvement of PDLP1 during *Hpa* infection, it is yet to establish whether the plasmodesmal immunity is involved. It has also been demonstrated that pathogen-associated molecular patterns (PAMPs), the fungal cell wall PAMP (chitin) or a 22-amino acid peptide of bacterial flagellin (flg22), are sufficient to trigger callose deposition at PD in Arabidopsis (Faulkner et al., 2013; Xu et al., 2017).

Recent findings began to reveal that pathogenic microbes utilize protein effectors to modulate the PD function in their hosts. Microbial effectors are known for altering plant cellular processes to suppress plant immunity (Cui et al., 2015; Buttner, 2016). The fungal pathogen *Fusarium oxysporum* effectors Avr2 and Six5 are localized to PD in *N. benthamiana* (Cao et al., 2018). The two PD-localized effectors form heterodimer and regulate the plasmodesmal function by allowing larger molecules to traffic through PD. The oomycete pathogen *Phytophthora brassicae* RxLR3 effector is localized to PD and physically associated with callose synthases, CalS1, CalS2, and CalS3. The expression of RxLR3 suppresses the function of the CalSs, inhibits the callose accumulation at PD, and promotes the PD-dependent movement of fluorescent molecules (Tomczynska et al., 2020). The bacterial pathogen *Pseudomonas syringae* pv. *tomato* DC3000 (*Pst* DC3000) delivers effector HopO1-1 to regulate the PD function by targeting PDLP5 and PDLP7 (Aung et al., 2020). Together, the reports showed that pathogenic microbes use effectors to target different PD regulators.

*Pst* DC3000 deploys 36 effectors into host cells through type III secretion system (Lindeberg et al., 2012; Wei et al., 2015). The regulation of PD by HopO1-1 prompted us to investigate whether bacterial effectors can move through PD in plants. In this study, we determined the PD-dependent movements of *Pst* DC3000 effectors in plants. We also explored how PDLP5, PDLP7, and flg22 affect the PD-dependent movement of bacterial effectors between plant cells.

## MATERIALS AND METHODS

### Plant growth conditions

*Nicotiana benthamiana* plants were grown at 22°C with 50% humidity and irradiated with 120 μmol m^-2^ s^-1^ white light for 14 h per day.

### Gene cloning and plasmid construction

To generate effector with 2 tandem repeats of YFP, the coding sequence of YFP without a stop codon was amplified from pGW2-YFP (Reumann et al., 2009). PCR products of effectors and YFP were fused together using an overlapping PCR method with Gateway-compatible primers as described previously (Aung et al., 2020). The stitched PCR fragments were cloned into pDONR 207 and a destination vector pGW2-YFP (Reumann et al., 2009) using a standard Gateway cloning system (Invitrogen). To construct a HF-mCherry construct, the coding sequence of mCherry was amplified from mCherry-pTA7002 (Fujioka et al., 2007) using Gateway-compatible primers. The PCR product was cloned into pDONR 207 and a destination vector pB7-HFN-stop (Lee et al., 2017). All primers used for cloning are listed in Supplemental Table 2.

### Agrobacterium-Mediated Transient Expression

*Agrobacterium tumefaciens* GV3101 harboring different expression constructs were cultured in a 30°C shaking incubator overnight. The overnight cultures were adjusted to a desired bacterial density using sterilize ddH2O. The bacterial solutions were infiltrated into fully expended leaves of 5-week-old *N. benthamina*. For subcellular localization analysis and immunoblot analysis a bacterial density of 0.1 (Aβoo) was used. Samples were collected 2-days after infiltration for confocal imaging or immunoblot assays.

### PD-Dependent Movement Assay

To determine the movement of effectors between plant cells, we adopted an established Agrobacterium-mediated protein movement assay (Brunkard et al., 2015). In brief, Agrobacteria harboring different expression cassettes of bacterial effector-fused with YFP were infiltrated into the fourth leaf of 5-week-old *N. benthamiana* at Aβoo 2×10^-4^. The expression of the fusion proteins was detected 2 days post infection using confocal microscopy. Plant cells with strong YFP signals were designated as the transformed plant cells. The movement of the effector fusion proteins was determined by the detection of YFP signals surrounding the transformed cells. The PD-dependent movement was quantified by scoring the movement efficiency of YFP. If YFP is only detected in the transformed cells, they are scored as 0. If YFP is detected in cells physically connecting the transformed cells, they are scored as 1. If YFP diffused beyond the first cell layer from the transformed cells, they are scored as ≥2. Over 100 transformation events were imaged across at least three biological replicates for all effectors except AvrE-YFP. Only 48 images were collected for AvrE-YFP from three biological replicates. The data were plotted with Excel. To compared the movement between effector-YFP and effector-2×YFP, the numbers of surrounding plant cells to the transformed cells containing YFP signals were counted. Over 100 images collected from three biological replicates were analyzed.

To determine the effect of PDLP5 and PDLP7 on the movement of a bacterial effector HopAF1, a mixture of PDLP5-HF (A_600_ 0.1), PDLP7-HF (A_600_ 0.1), or HF-mCherry (A_600_ 0.1) with HopAF1-YFP (A_600_ 2×10^-4^) was co-infiltrated into fully expanded leaves of 5-week-old *N. benthamiana*. The movement of HopAF1-YFP was determined by the detection of YFP signals surrounding the transformed cells as described above. The numbers of surrounding plant cells to the transformed cells containing YFP signals were counted and compared. Over 100 images collected from three biological replicates were analyzed.

To determine the effect of flg22 on the PD-dependent movement of a bacterial effector HopAF1, 0.1μM of flg22 was infiltrated into fully expanded leaves of 5-week-old *N. benthamiana*. For a mock treatment, ddH20 was infiltrated. 24 hours after the treatment, Agrobacteria harboring HopAF1-YFP were infiltrated into the mock- or flg22-treated leaves. Over 100 images collected from three biological replicates were analyzed.

### Plasmodesmal callose staining assay

The fourth leaf of *N. benthamiana* was infiltrated with ddH2O (mock) or 0.1μM of flg22 for 24 hours. Aniline blue (0.01% in 1 ×PBS buffer, pH 7.4) was infiltrated into the treated area to image callose accumulation at PD as previously described (Xu et al., 2017). To determine the role of PDLP5 and PDLP7 in callose accumulation, Agrobacteria harboring *35S::PDLP5-HF, 35S::PDLP7-HF*, and *35S::HF-YFP* (mock) were infiltrated into the fourth leaf of *N. benthamiana*. Aniline blue was infiltrated into the bacterial infected area 48 hours post infection for imaging callose accumulation at PD as mentioned above. Plasmodesmal callose deposits were imaged using confocal microscopy 15 min after dye infiltration. 10 images were collected from each sample. Aniline blue stained callose was quantified using the Macro feature of FIJI for large scale data analysis. In brief, images were first converted from lsm to tif then to 8-bit image files. RenyiEntropy white method was used to set Auto Threshold creating black and white images highlighting callose. Particle Analysis tool was used to outline each aniline blue-stained callose and ascribe a quantitative numerical value in μm^2^. Exclusion setting of 0.10-20 μm^2^ and a circularity of 0.30-1.00 were used to isolate callose excluding any non callose related fluorescence. 10 images were collected from each treatment. Data from 10 images from an experiment were pooled and plotted as mentioned below.

### Statistical Analysis

All presented experiments were performed at least three independent times. The pooling of data from different biological replicates for different experiments is indicated in each section. Violin box plots were created with an online software (https://huygens.science.uva.nl/PlotsOfData/). Mann-Whitney *U* Test (https://www.socscistatistics.com/tests/mannwhitney/default2.aspx) was performed for testing statistical significance of differences.

### Immunoblot analyses

*N. benthamiana* leaves were frozen with liquid nitrogen and homogenized with 1600 miniG (SPEX). Protein extraction buffer (60 mM Tris-HCl [pH 8.8], 2% [v/v] glycerol, 0.13 mM EDTA [pH 8.0], and 1 × protease inhibitor cocktail complete from Roche) was added to the homogenized tissues (100 μl/10 mg). The samples were vortexed for 30 sec, heated at 70°C for 10 min, and centrifuged at 13,000*g* for 5 min at room temperature. The supernatants were then transferred to new tubes. For SDS-PAGE analysis, 10 μl of the extract in 1x Laemmli sample buffer (Bio-Rad) was separated on 4–15% Mini-PROTEAN TGX precast protein gel (Bio-Rad). The separated proteins were transferred to a polyvinylidene fluoride membrane (Bio-Rad) using a Trans-Blot Turbo Transfer System RTA transfer kit following the manufacturer’s instructions (Bio-Rad). The membrane was incubated in a blocking buffer (3% [v/v] BSA, 50 mM Tris base, 150 mM NaCl, 0.05% [v/v] Tween 20 [pH 8.0]) at room temperature for 1 hr, then incubated overnight with an antibody prepared in the blocking buffer at 4°C overnight. The antibodies used are as follows: 1:20,000 a-GFP (Abcam catalog No. ab290), 1:10,000 a-cMyc (Abcam catalog No. ab9106), and 1:10,000 a-Flag-HRP (Sigma-Aldrich catalog No. A8592). The probed membranes were washed three times with 1× TBST (50 mM Tris base, 150 mM NaCl, and 0.05% [v/v] Tween 20, pH 8.0) for 5 min before being incubated with a secondary antibody at room temperature for 1 hr except for a-Flag-HRP. The secondary antibodies used were 1:20,000 goat anti-rabbit IgG (Thermo Fisher Scientific catalog No. 31,460). Finally, the membranes were washed four times with 1 × TBST for 10 min before the signals were visualized with SuperSignal West Dura Extended Duration Substrate (Pierce Biotechnology).

## RESULTS

### Bacterial effectors traffic between plant cells

To determine whether effectors can move from the infected cells to the surrounding plant cells, we monitored the movement of yellow fluorescent protein (YFP) tagged *Pst* DC3000 effector-YFPs (Aung et al., 2020). 29 *Pst* DC3000 effector-YFPs were transiently expressed in *N. benthamiana* using an Agrobacterium-mediated transient expression method. Transient expression of some effectors leads to plant cell death, whereas the expression of a few effectors could not be detected. We thus selected 17 effectors to further investigate their movement between plant cells (Figure 1 and Supplemental Figure 1). We adopted an established Agrobacterium-mediated protein movement assay to test whether effectors are able to move between plant cells (Brunkard et al., 2015). A relatively low inoculum (A_600_ 2×10^-4^) of *Agrobacterium tumefaciens* harboring the plasmid of interests was infiltrated into *N. benthamiana*, resulting in a few transformed cells per cm^2^. The transformed plant cells exhibit strong fluorescence signals. If the fusion proteins move beyond initially transformed cells, weak fluorescence signals are observed in the cells surrounding the transformed cells. Using confocal microscopy, we observed the movement of 16 bacterial effectors between plant cells (Figure 1). Over 50% of transformation events leads to the intercellular movement of HopK1-YFP, HopF2-YFP, HopH1-YFP, and HopAF1-YFP. Among them, HopAF1-YFP shows the most effective movement. Over 20% of transformation events results in the trafficking of HopAF1-YFP to two or more than two cell layers from the transformed cells (Figure 1b). For the majority of mobile effectors, around 20-30% of transformed cells exhibits the movement beyond initially transformed cells (Figure 1b). We then conducted immunoblot analysis to confirm the expression of full-length fusion proteins. Total proteins were extracted from *N. benthamiana* leaves transiently expressing the effector fusion proteins. The expression of the fusion proteins was detected using a GFP antibody. As shown in Supplemental Figure 2a, we detected a major band for most effectors at higher molecular weight, suggesting that fluorescence signals detected in Figure 1a are emitted from full-length effector fusion proteins. It is noted that most fusion proteins migrated slower than expected (Supplemental Figure 2a and Supplemental Table 1). Together, the findings suggest that bacterial effectors are able to move beyond initially transformed cells.

**Figure 1.**
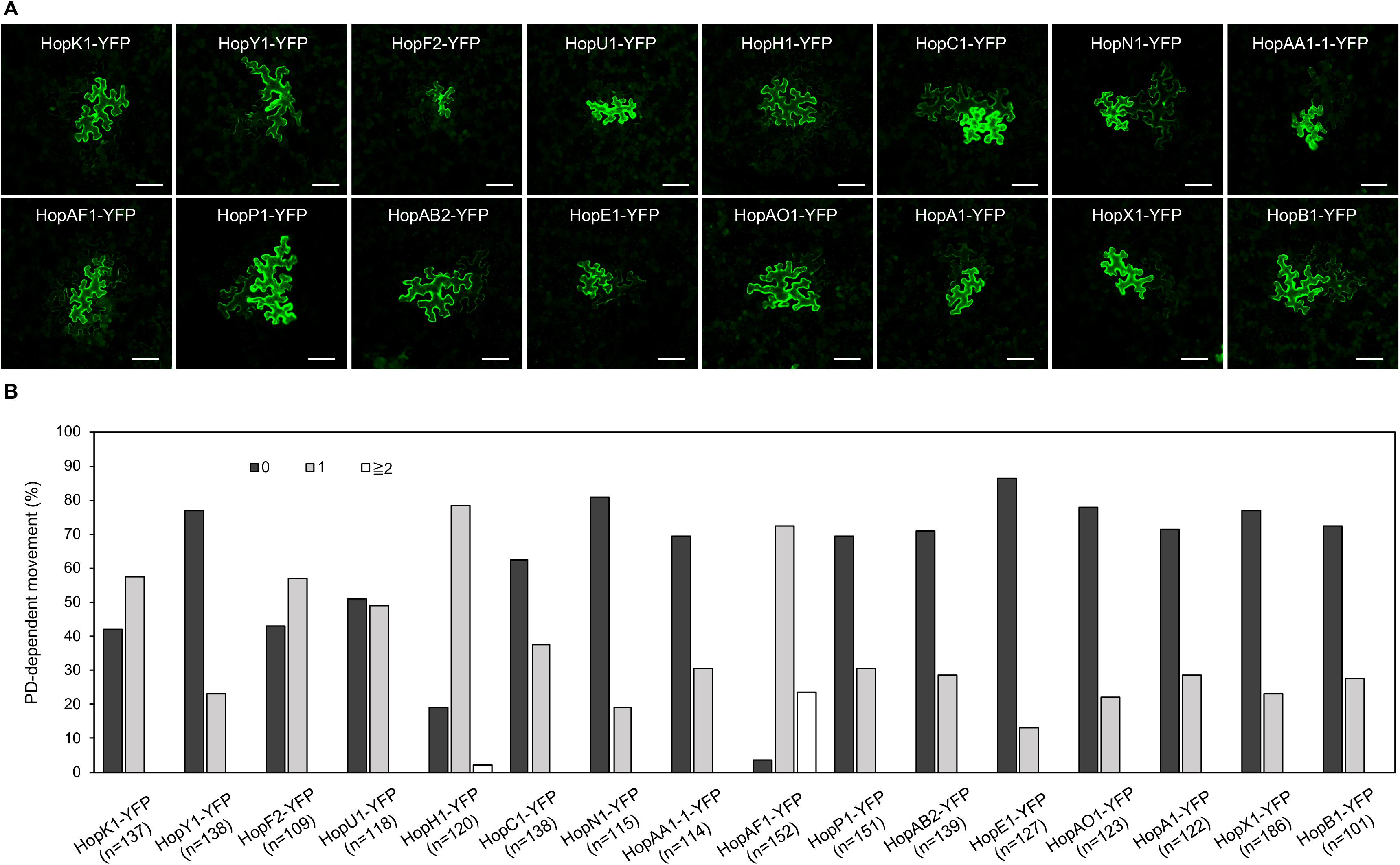
The movement of bacterial effectors between plant cells. **(A)** Confocal images show the diffusion of effector-YFP fusion proteins. Images were taken from epidermis of *N. benthamiana* leaves. An agrobacterium-mediated protein movement assay was conducted to determine the movement of effector-YFP fusion proteins in plants. The transformed plant cell exhibits strong yellow fluorescent (YFP) signals. The movement of the fusion proteins is determined by the detection of YFP signals in cells surrounding the transformed cell. Scale bars = 100 μm. **(B)** Quantitative data show the percentage of transformation events resulting in no diffusion (0), one cell layer diffusion (1), and two or more than 2 cell layers diffusion (≧2). The data shown here are pooled from at least three biological replicates. The number of transformation events analyzed is indicated (n).

### The movement of effectors largely depends on their molecular weights

Predicted molecular weights of most effector-YFP fusion proteins ranged between 50-80 kDa, whereas a few effectors like HopR1 and AvrE weight over 200 kDa (Supplemental Table 1). The majority of the mobile effector-YFPs shown in Figure 1 weights below 70 kDa, expect HopAA1-1-YFP (77.6 kDa; Supplemental Table 1). Among the tested effectors, AvrE-YFP doesn’t move from transformed cells to the neighboring cells (Figure 2A–2B). As *AvrE-YFP* encodes a protein with the molecular weight of ~222 kDa, the large molecule might impede the PD-dependent movement of the effector. It is noted that the expression of AvrE-YFP using higher Agrobacterium inoculum (A_600_ 0.1) leads to cell death, whereas lower Agrobacterium inoculum (A_600_ 2×10^-4^) allows us to detect the expression of the fusion proteins two days after infiltration.

**Figure 2.**
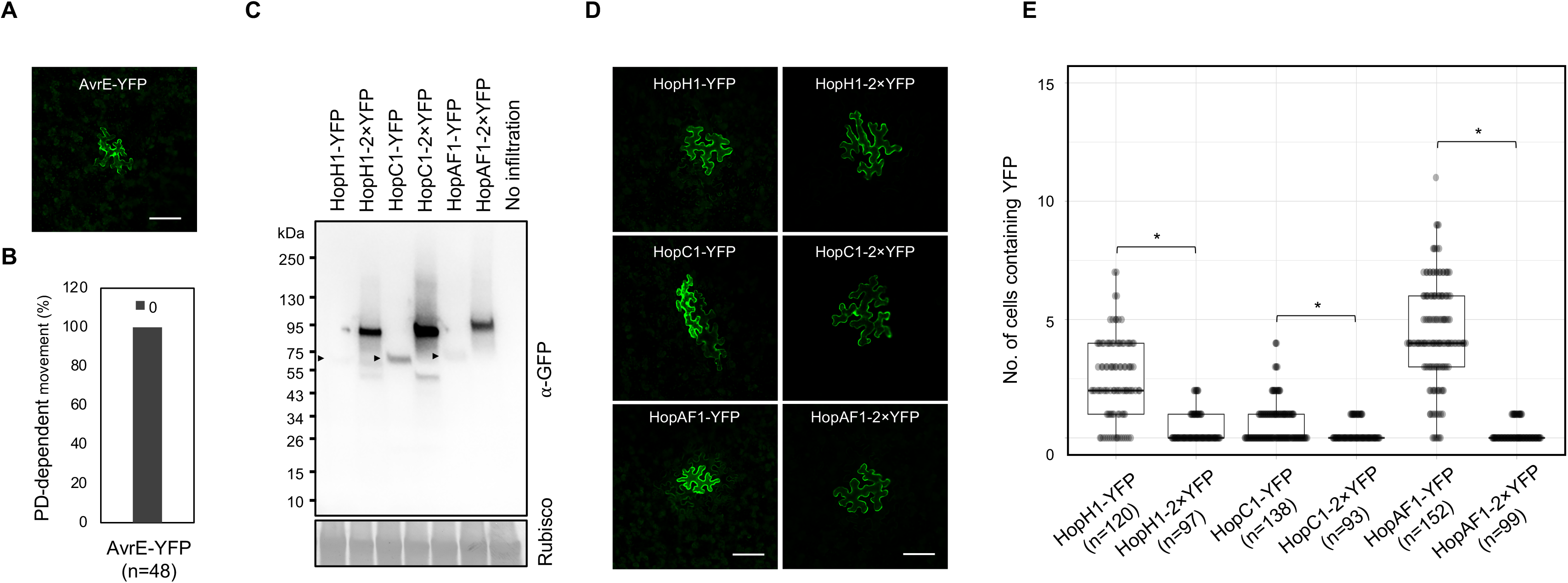
The movement of effectors is largely affected by their sizes. **(A)** Confocal image shows the expression of AvrE-YFP. Scale bars = 100 μm. **(B)** Quantitative data show that AvrE-YFP cannot move through PD. 0: no diffusion. The number of transformation events analyzed is indicated (n). **(C)** Detection of full-length effector fusion proteins. Effector-YFPs and effector-2×YFPs were transiently expressed in *N. benthamiana*. The samples were then subjected to immunoblot analysis using an anti-GFP antibody. Rubisco is served as a loading control. Arrow heads indicate the expression of effector-YFPs. **(D)** Confocal images show the diffusion of effector-YFP fusion proteins. Images were taken from epidermis of *N. benthamiana* transiently expressing different fusion proteins. Scale bars = 100 μm. **(E)** Quantitative data present the PD-dependent movement of effector-YFP fusion proteins. Mann-Whitney *U* Test was used to analyze the data. The p-value is <0.0001 for HopH1-YFP vs. HopH1-2×YFP, <.00094 for HopC1-YFP and HopC1-2×YFP, and <0.00001 for HopAF1-YFP and HopAF1-2×YFP (*). The number of transformation events analyzed is indicated (n).

As we hypothesized that effectors move intercellularly through PD, we next examined whether the molecular weights of the mobile effectors affects their movement. We thus constructed HopH1, HopC1, and HopAF1 with two tandem repeats of YFP, yielding HopH1-2×YFP, HopC1-2×YFP, and HopAF1-2×YFP. We first determined the molecular weights of the fusion proteins by transiently expressing them in *N. benthamiana* leaves using an Agrobacterium-mediated approach. We then detected the expression of the fusion proteins using a GFP antibody. Compared to 1 ×YFP fusion, 2×YFP fusion of the effectors increases the molecular weight by ~26 kDa (Figure 2C). To investigate the PD-dependent movement of 2×YFP fusion proteins, we determined the movement of HopH1-2×YFP, HopC1-2×YFP, and HopAF1-2×YFP in *N. benthamiana* leaves as mentioned above. Compared to the diffusion of 1×YFP fusion proteins, the movement of HopH1-2×YFP, HopC1-2×YFP, and HopAF1-2×YFP beyond initially transformed cells is drastically reduced (Figure 2D and 2E). Together, the findings support that the bacterial effectors move between plant cells through PD.

### The expression of PDLP5 and PDLP7 suppresses the PD-dependent movement of HopAF1

Altered expression of PDLPs has been shown to impact the PD function. To further support that the intercellular movement of effectors depends on PD, we investigated whether the expression of PDLP affects the movement of bacterial effectors. PDLP5 has been shown to affect callose deposition at PD and alter the movements of GFP molecules between cells in Arabidopsis; however, it’s unknown whether PDLPs regulates callose accumulation at PD when transiently expressed in *N. benthamiana*. To this end, we detected callose accumulation at PD in *N. benthamiana* after PDLP5 or PDLP7 was transiently expressed. PDLP5 and PDLP7 were selected due to their role in bacterial immunity (Lee et al., 2011; Aung et al., 2020). We first detected the expression of HF-YFP (mock), PDLP5-HF, and PDLP7-HF using immunoblot analysis (Supplemental Figure 2B). The leaf transiently expressing the fusion proteins was stained with aniline blue to detect callose accumulated at PD. Similar to Arabidopsis transgenic plants overexpressing PDLP5 (Lee et al., 2011), transient expression of Arabidopsis PDLP5 is sufficient to increase callose accumulation at PD compared to that of mock treatment (Figure 3A–3B). While PDLP5 has been previously shown to regulate callose homeostasis, whether PDLP7 has similar roles in callose accumulation has not been determined. Here we demonstrated that transient expression of Arabidopsis PDLP7 also leads to higher accumulation of callose at PD in *N. benthaimana* (Figure 3A–3B). Together, the findings showed that transient overexpression of the PDLP5 and PDLP7 could increase the callose accumulation at PD in *N. benthamiana*.

**Figure 3.**
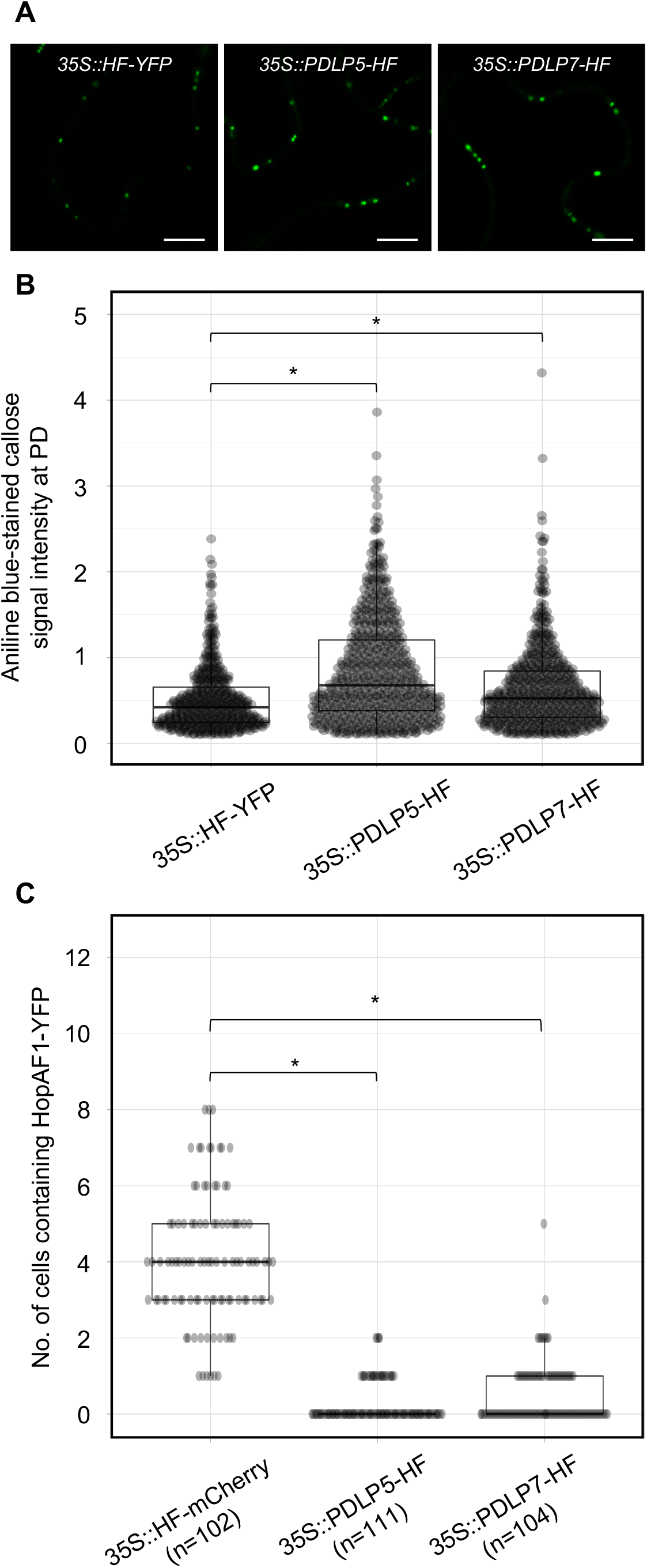
Expression of PDLP5 and PDLP7 suppresses the PD-dependent movement of HopAF1. **(A)** Expression of PDLPs and HopO1-1 affects callose accumulation at PD. *N. benthamiana* leaves were infiltrated with Agrobacteria harboring *35S::HF-YFP, 35S::PDLP5-HF*, and *35S::PDLP7-HF*. Infiltrated leaves were stained with 0.01% aniline blue and imaged with confocal microscopy. Scale bars = 10 μm. **(B)** Quantitative data present the accumulation of callose at PD. Mann-Whitney *U* Test was used to analyze the data. The p-value is <0.0001 (*). **(C)** Quantitative data present the PD-dependent movement of HopAF1-YFP when co-expressed with *35S::HF-mCherry, 35S::PDLP5-HF*, and *35S::PDLP7-HF*. Mann-Whitney *U*Test was used to analyze the data. The p-value is <0.00001 for both *35S::PDLP5-HF* and *35S::PDLP7-HF* (*). The number of transformation events analyzed is indicated (n).

Callose accumulation at PD is negatively associated with PD-dependent movement of molecules between plant cells (De Storme and Geelen, 2014; Amsbury et al., 2017; Wu et al., 2018). To further support that bacterial effectors move through PD, we investigated whether PDLP-mediated PD closure would suppress the movement of a bacterial effector HopAF1. Among the mobile effectors, HopAF1 was chosen in this assay because of its highest PD-dependent movement in plants (Figure 1). Relatively lower inoculum (A_600_ 2×10^-4^) of Agrobacteria harboring *35S::HopAF1-YFP* was mixed with higher inoculum (A_600_ 0.1) of Agrobacteria harboring *35S::HF-mCherry* (mock), *35S::PDLP5-HF*, or *35S::PDLP7-HF* and infiltrated into *N. benthamiana* leaves. The expression of the fusion proteins was determined using immunoblot analysis (Supplemental Figure 2C). The movement of HopAF1-YFP was determined two days after the Agrobacterium infiltration using confocal microscopy. As shown in Figure 3C, the expression of PDLP5 and PDLP7 drastically reduced the intercellular movement of HopAF1-YFP beyond the transformed cells. Together, the findings suggest that the expression of PDLP5 and PDLP7 affects the PD-dependent movement of a bacterial effector.

### flg22 inhibits the movement of a bacterial effector

In addition to PDLP expression, flg22 has been also reported to induce callose deposition at PD and reduce the PD-dependent molecular fluxes between cells in Arabidopsis (Faulkner et al., 2013; Xu et al., 2017). To determine the effect of flg22 on callose accumulation at PD in *N. benthamiana*, we treated a fully expended leaf of *N. benthamina* with 0.1 μM flg22. Callose deposition at PD was examined 24 hours after the infiltration. flg22-treated leaf, compared to mock-treated leaf (infiltrated with ddH2O), exhibits higher accumulation of callose at PD (Figure 4A–4B). We next determined whether flg22 treatment suppresses the movement of HopAF1 using an Agrobacterium-mediated protein movement assay mentioned above. The expression of HopAF1-YFP was detected using immunoblot analysis (Supplemental Figure 2D). In line with the callose accumulation at PD, flg22-treatment inhibits the PD-dependent movement of HopAF1 (Figure 4C).

**Figure 4.**
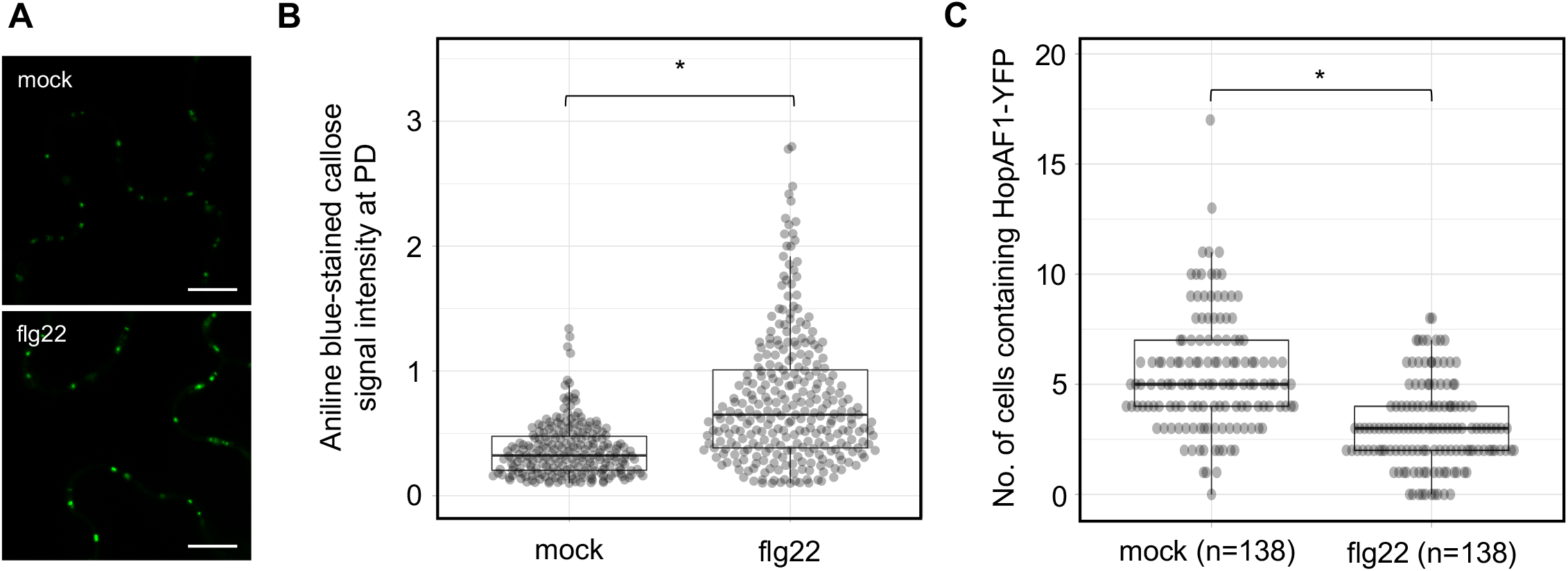
PTI-induced callose accumulation at PD reduces the movement of effectors. **(A)** flg22 induces callose accumulation at PD. *N. benthaimiana* leaves were infiltrated with 0.1 nM of flg22 or ddH2O (mock). Infiltrated leaves were stained with 0.01% aniline blue and imaged with confocal microscopy. Scale bars = 10 μm. **(B)** Quantitative data present the accumulation of callose at PD. Mann-Whitney *U* Test was used to analyze the data. The p-value is <0.00001 (*). **(C)** flg22 treatment suppresses the PD-dependent movement of HopAF1. *N. benthaimiana* leaves were pretreated with 0.1 nM of flg22 or ddH2O (mock) for 24 hr. Agrobacteria harboring *35S::HopAF1-YFP* were later infiltrated into the pretreated leaves. The PD-dependent movement of HopAF1-YFP was examined 48 hr post Agrobacterium infiltration using confocal microscopy. The data shown here were collected from four biological repeats. Mann-Whitney *U* Test was used to analyze the data. The p-value is <0.00001 (*). The number of transformation events analyzed is indicated (n).

## DISCUSSION

The PD-dependent movement of fungal effectors (Khang et al., 2010) and an oomycete effector (Khang et al., 2010; Cao et al., 2018; Tomczynska et al., 2020) have been reported; however, it’s unclear whether bacterial effectors move between plant cells or not. Empirical evidence from this work showed that at least 16 *Pst* DC3000 effectors move between plant cells through PD. We established that the movement of the effectors is dependent on PD from the following findings: (1) the effector-YFP fusion proteins can move from the transformed cells to the adjoining plant cells (Figure 1), (2) the movement of the effectors is largely dependent on their molecular weights (Figure 2), and (3) PDLP5- and PDLP7-induced callose accumulation at PD inhibits the movement of a bacterial effector HopAF1 (Figure 3).

Although the movement of 16 effectors is reported here, it’s plausible that more *Pst* DC3000 effectors are able to move between plants cells. The following reasons might account for the underestimation of the PD-dependent movement of bacterial effectors: (1) the YFP fusion of effectors increases their molecular weights and suppresses their PD-dependent movement, (2) transiently overexpressing individual effector induces cell death in *N. benthamiana* thus preventing the visualization of the effectors, and (3) the expression level of some effectors is under the detectable threshold using confocal microscopy.

It’s well established that the molecular weight of proteins affects their movement between plant cells through PD (Kim et al., 2005; Aung et al., 2020). Among the effectors we investigated, we did not observe the movement of AvrE-YFP (Figure 2A–2B), which has an expected molecular weight of ~220 kDa. Also, the tandem fusion of YFP to HopH1-YFP, HopC1-YFP, and HopAF1-YFP (yielding HopH1-2×YFP, HopC1-2×YFP, and HopAF1-2×YFP) drastically suppresses the PD-dependent movement (Figure 2D–2E). The addition of another YFP increases the molecular weight of the fusion proteins by ~26 kDa (Figure 2C). It is also possible that the tandem fusion of YFP affects the tertiary structure of the fusion proteins, impeding the PD-dependent movement.

Many mobile effectors were detected both in the nucleus and cytoplasm (Supplemental Figure 1); however, we observed the PD-dependent movement of a few PM localized effectors. For example, HopAF1 was previously demonstrated to associate with the PM (Washington et al., 2016). We also observed the PM association of HopAF1 (Supplemental Figure 1). Interestingly, HopAF1 is the most mobile effector among the 16 effectors reported here (Figure 1B). Similarly, another PM-localized effector HopA1 is also mobile between plant cells (Figure 1). Together, the findings suggest that the membrane association of effectors does not inhibit the PD-dependent movement. It is unclear whether mitochondrial, chloroplast, or the endoplasmic reticulum association of effectors affects the PD-dependent movement in plants.

In Arabidopsis, the expression of PDLP5 is positively correlated with the accumulation of callose at PD (Lee et al., 2011). Here, we reported that the transient overexpression of Arabidopsis PDLP5 in *N. benthamiana* increases the accumulation of callose at PD (Figure 3A–3B). The finding is supported by a recent report that the transient overexpression of PDLP5 suppresses the PD-dependent movement of mCherry (Wang et al., 2020). Similar to PDLP5, the transient expression of Arabidopsis PDLP7 also increases callose accumulation at PD in *N. benthamiana* (Figure 3A–3B). Among different PDLP members, only the expression of PDLP5 transcripts and proteins is upregulated by bacterial infections and a defense hormone SA treatment (Lee et al., 2011). PDLP7 proteins are destabilized by *P. syringae* infection in a bacterial effector HopO1-1-dependent manner (Aung et al., 2020). Given that HopO1-1 physically associates with and destabilizes PDLP5 and PDLP7, the effector might target the PDLPs to suppress plasmodesmal immunity. The targeting of the PDLPs might play critical role in facilitating the PD-dependent movement of bacterial effectors from the infected cells to the adjoining non-infected cells through PD. In line with the notion, we observed that the transient overexpression of PDLP5 and PDLP7 significantly suppresses the PD-dependent movement of a highly mobile bacterial effector HopAF1.

PAMP-triggered callose accumulation at PD suggests that the plasmodesmal closure is a part of pattern-triggered immunity (PTI). PTI is considered the first line of plant immune responses during microbial infection (Boller and He, 2009). It is plausible that plants induce the plasmodesmal closure to limit the spread of microbial molecules from the infected cells to the surrounding plant cells. In line with the statement, flg22 treatment suppresses the PD-dependent movement of a bacterial effector HopAF1 (Figure 4C). It is postulated that the PTI-triggered callose accumulation at PD generally suppresses the PD-dependent movement of most effectors. Recent report showed that the expression of a bacterial effector HopO1-1 facilitates the PD-dependent movement of YFP molecules (Aung et al., 2020). As HopO1-1 targets and destabilizes PDLP5 and PDLP7, it is highly plausible that HopO1-1 functions to overcome the plasmodesmal immunity. We thus hypothesize that HopO1-1 might facilitate the PD-dependent movement of bacterial effectors to the surrounding plant cells. The hypothesis is supported by a recent report that the PD-dependent cell-to-cell movement of *Fusarium oxysporum* effector Avr2-GFP requires Aix5 (Cao et al., 2018). Further studies will reveal the role of HopO1-1 in modulating the PD-dependent movement of bacterial effectors.

As effector proteins are involved in suppressing plant immunity to benefit the microbes, the mobile effectors might play crucial roles in inhibiting non-cell-autonomous plant immunity. Successful suppression of both cell-autonomous and non-cell-autonomous plant immunity might be critical for pathogenic microbes to colonize and spread from the initial infection sites. Understanding the function of mobile effectors will allow us to better understand how pathogenic microbes regulate cellular processes in infected plant cells and the surrounding plant cells. This report also demonstrates that Agrobacterium-mediated protein movement assay using *N. benthamiana* is a powerful experimental system to determine the PD-dependent movement of microbial effectors. The system has great potential in directly visualizing how the mobile effectors affect plant immune responses. Identification and characterization of robust plant immune response markers will allow us to investigate the functions of mobile effectors in modulating cell-autonomous and non-cell-autonomous immune responses *in planta*.

## Data Availability Statement

All data generated for this study are included in the article/**Supplementary Material**, further inquiries can be directed to the corresponding author.

## Author Contributions

KA designed the research and conducted the experiments shown in Figure 3C, Supplemental Figure 1, Supplemental Figure 2, and wrote the manuscript with input from all authors. ZPL performed the experiments shown in Figure 1, and Figure 2. HV conducted the experiments shown in Figure 3A, Figure 3B, and Figure 4. YC conducted the experiments shown in Figure 2D and Figure 2E. SLL contributed to Figure 2C and Supplemental Figure 2.

## Acknowledgements

We would like to thank Dr. Sheng Yang He for his generosity. Most constructs used in this study were cloned by KA when he was a postdoc in the He laboratory at Michigan State University. We also would like to thank Tyler Weide for growing *N. benthaimana*.

## CONTRIBUTION TO THE FIELD STATEMENT

Multicellular organisms are armed with sophisticated immune systems to defend against pathogens. Pathogenic microbes, on the other hand, could deploy effector proteins into their host cells to suppress the immune systems. Tremendous progress has been made toward understanding host immune responses at pathogen-infected cells, cell-autonomous immunity; however, it is largely unknown how pathogens overcome host immunity at the neighboring host cells surrounding the pathogen-infected cells. Recent findings revealed that microbial effectors target plant cell-to-cell communication channels, plasmodesmata (PD). In addition, a few fungal effector proteins have been reported to move beyond initial infected cells through PD. These findings support that pathogens suppress plant immune responses in the surrounding non-infected plant cells, non-cell-autonomous immunity. In this study, we demonstrated the PD-dependent movement of the bacterial pathogen *Pseudomonas syringae* pv. *tomato* DC3000 effectors in plants using an Agrobacterium-mediated protein movement assay in *Nicotiana benthamiana*. The mobile effectors identified here represent as ideal tools to further investigate how pathogens manipulate cell-autonomous and non-cell-autonomous immunity in live tissues. Further studies will reveal the dynamic interactions between plants and pathogens as well as the crucial role of PD during defense.

## Supplementary Material

**Supplemental Figure 1.** Expression and subcellular localization of bacterial effector proteins transiently expressed in *N. benthamiana*.

**Supplemental Figure 2.** Immunoblot analysis detects the expression of different fusion proteins in *N. benthamiana*.

**Supplemental Table 1.** Estimated molecular weights of effector-YFP fusion proteins.

**Supplemental Table 2.** Primers used for cloning in this study.

**Supplemental Figure 1. Expression and subcellular localization of bacterial effector proteins transiently expressed in *N. benthamiana*.** Images were taken with confocal microscopy. Scale bars = 10 μm.

**Supplemental Figure 2. Immunoblot analysis detects the expression of different fusion proteins in *N. benthamiana*.**

**(A)** A GFP antibody was used to detect the expression of full-length effector-YFP fusion proteins.

**(B)** A Flag antibody was used to detect the expression of HF-YFP, PDLP5-HF, and PDLP7-HF fusion proteins.

**(C)** A Flag antibody and GFP antibody were used to detect the expression of HF-YFP, PDLP5-HF, and PDLP7-HF when co-infiltrated with HopAF1-YFP. -: without expressing HF fusion proteins.

**(D)** A GFP antibody was used to detect the expression of HopAF1-YFP fusion protein with (+) or without (-) flg22 treatment.

