## Supplemental Figures for "Plasmodesmata-dependent intercellular movement of bacterial effectors"

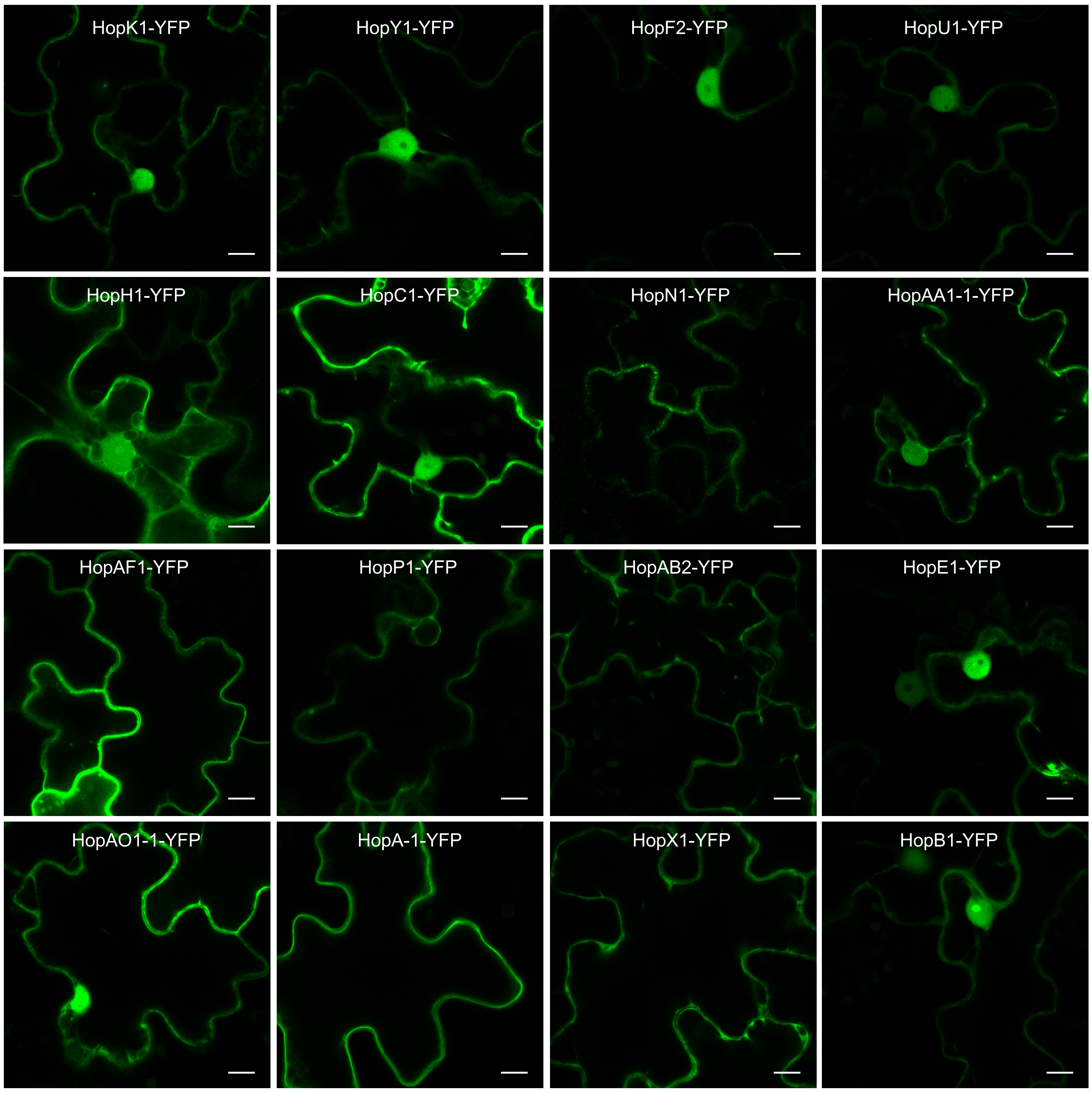


**Supplemental Figure 1. Expression and subcellular localization of bacterial effector proteins transiently expressed in *N. benthamiana*.** Images were taken with confocal microscopy. Scale bars = 10 μm.

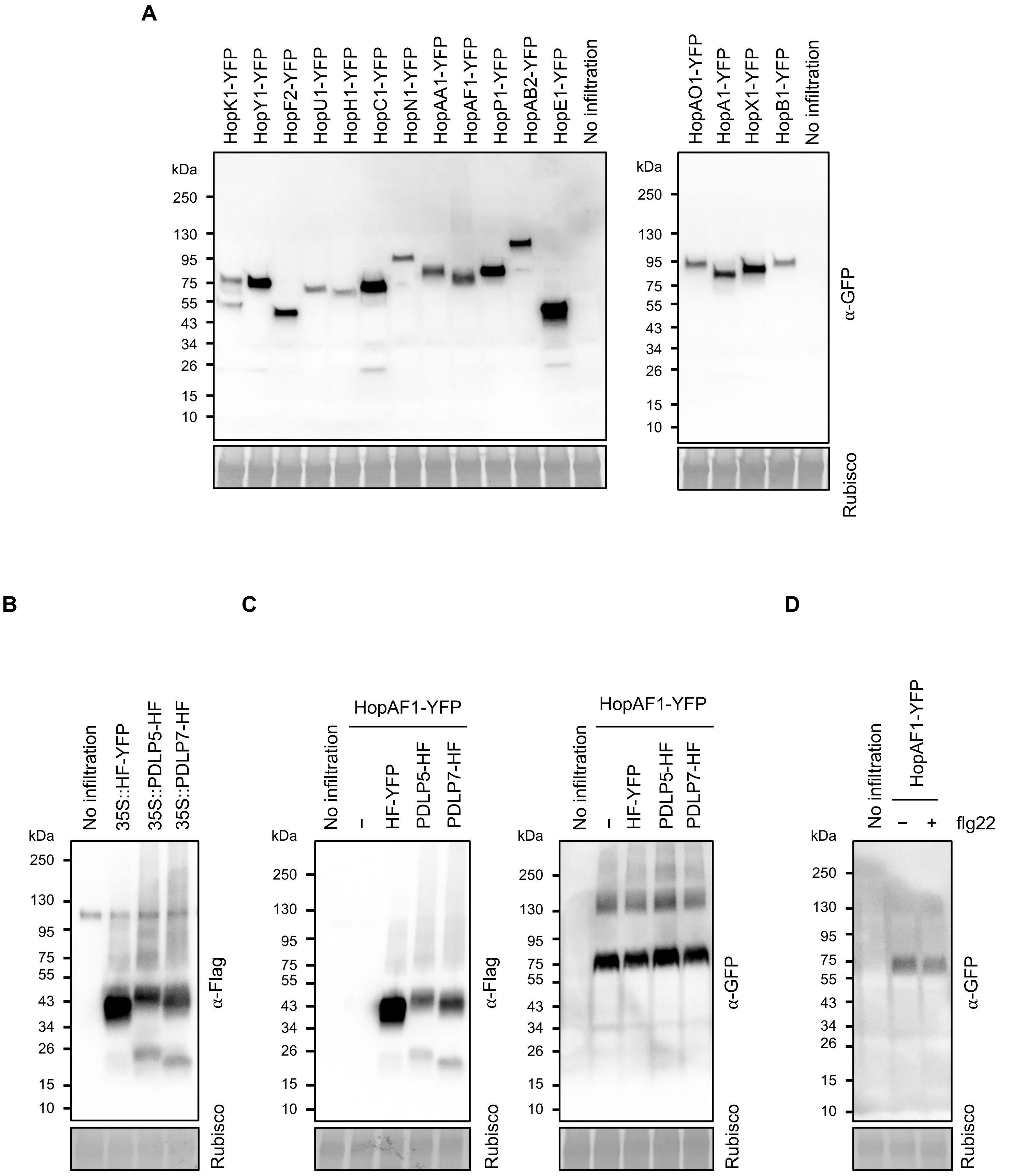


**Supplemental Figure 2. Immunoblot analysis detects the expression of different fusion proteins in *N. benthamiana*.**
(A) A GFP antibody was used to detect the expression of full-length effector-YFP fusion proteins.
(B) A Flag antibody was used to detect the expression of HF-YFP, PDLP5-HF, and PDLP7-HF fusion proteins.
(C) A Flag antibody and GFP antibody were used to detect the expression of HF-YFP, PDLP5-HF, and PDLP7-HF when co-infiltrated with HopAF1-YFP. −: without expressing HF fusion proteins.
(D) A GFP antibody was used to detect the expression of HopAF1-YFP fusion protein with (+) or without (−) flg22 treatment. Three biological replicates were performed and exhibited similar results.
