## Supplemental Table 1 for "Plasmodesmata-dependent intercellular movement of bacterial effectors"

Supplemental Table 1. Estimated molecular weights of effector-YFP fusion proteins.

| Fusion protein | Molecular weight (kDa) |
| --- | --- |
| HopK1-YFP | 63.4 |
| HopY1-YFP | 57.9 |
| HopF2-YFP | 49.3 |
| HopU1-YFP | 56.9 |
| HopH1-YFP | 51.2 |
| HopC1-YFP | 55.8 |
| HopN1-YFP | 65.4 |
| HopAA1-1-YFP | 77.6 |
| AvrE1-YFP | 222 |
| HopAF1-YFP | 57.8 |
| HopP1-YFP | 59.3 |
| HopAB2-YFP | 86.3 |
| HopE1-YFP | 50.9 |
| HopAA1-2-YFP | 78 |
| HopAO1-YFP | 78.3 |
| HopI1-YFP | 79.5 |
| HopA1-YFP | 68.9 |
| HopX1-YFP | 68.3 |
