## Supplemental Table 2 for "Plasmodesmata-dependent intercellular movement of bacterial effectors"

Supplemental Table 2. Primers used for cloning in this study.

| Primer | Sequence | For cloning |
| --- | --- | --- |
| HopH1-attB1 | ggggacaagtttgtacaaaaaagcaggcttcatgatcactccgtctcgatatc | HopH1-2×YFP |
| HopH1-YFP-F | gtacagggcacatcaaatggtgagcaagggcg | HopH1-2×YFP |
| HopH1-YFP-R | cgcccttgctcaccatttgatgtgccctgtac | HopH1-2×YFP |
| EYFP-attB2 | ggggaccactttgtacaagaaagctgggtcttacttgtacagctcgtcc | HopH1-2×YFP |
| HopC1-attB1 | ggggacaagtttgtacaaaaaagcaggcttcatgacaatcgtgtctggac | HopC1-2×YFP |
| HopC1-YFP-F | gtattcgcttcaaaaatacacatggtgagcaagggc | HopC1-2×YFP |
| HopC1-YFP-R | gcccttgctcaccatgtgtatttttgaagcgaatac | HopC1-2×YFP |
| EYFP-attB2 | ggggaccactttgtacaagaaagctgggtcttacttgtacagctcgtcc | HopC1-2×YFP |
| HopAF1-attB1 | ggggacaagtttgtacaaaaaagcaggcttcatggggctatgtatttcaaaac | HopAF1-2×YFP |
| HopAF1-YFP-F | catctggtcgcacaaatggtgagcaagggcg | HopAF1-2×YFP |
| HopAF1-YFP-R | cgcccttgctcaccatttgtgcgaccagatg | HopAF1-2×YFP |
| EYFP-attB2 | ggggaccactttgtacaagaaagctgggtcttacttgtacagctcgtcc | HopAF1-2×YFP |
| mCherry-attB1 | ggggacaagtttgtacaaaaaagcaggcttcatggtgagcaagggcg | HF-mCherry |
| mCherry-attB2 | ggggaccactttgtacaagaaagctgggtcttacttgtacagctcgtcc | HF-mCherry |
